# Toxoflavin secreted by *Pseudomonas alcaliphila* inhibits growth of *Legionella pneumophila* and its host *Vermamoeba vermiformis*

**DOI:** 10.1101/2022.01.08.475489

**Authors:** Sebastien P. Faucher, Sara Matthews, Arvin Nickzad, Passoret Vounba, Deeksha Shetty, Émilie Bédard, Michele Prévost, Eric Déziel, Kiran Paranjape

**Affiliations:** Department of Natural Resource Sciences, Faculty of Agricultural and Environmental Sciences, McGill University, Sainte-Anne-de-Bellevue, QC, Canada; INRS–Centre Armand-Frappier Santé Biotechnologie, Laval, QC, Canada; Department of Civil Engineering, Polytechnique Montréal, Montréal, Québec, Canada; Department of Medical Biochemistry and Microbiology, Uppsala University, Uppsala, Sweden

**Keywords:** Toxoflavin, *Legionella*, *Pseudomonas*, water, biofilm

## Abstract

*Legionella pneumophila* is a natural inhabitant of water systems. From there, it can be transmitted to humans by aerosolization resulting in severe pneumonia. Most large outbreaks are caused by cooling towers contaminated with *L. pneumophila*. The resident microbiota of the cooling tower is a key determinant for the colonization and growth of *L. pneumophila*. The genus *Pseudomonas* correlates negatively with the presence of *L. pneumophila*, but it is not clear which species is responsible. Therefore, we identified the *Pseudomonas* species inhabiting 14 cooling towers using a *Pseudomonas*-specific 16S rRNA amplicon sequencing strategy. Cooling towers free of *L. pneumophila* contained a high relative abundance of members from the *Pseudomonas alcaliphila/oleovorans* phylogenetic cluster. *In vitro*, *P. alcaliphila* JCM 10630 inhibited the growth of *L. pneumophila* on agar plates. Analysis of the *P. alcaliphila* genome revealed the presence of a genes cluster predicted to produce toxoflavin. *L. pneumophila* growth was inhibited by pure toxoflavin and by extract from *P. alcaliphila* culture found to contain toxoflavin by LC-ESI-MS. In addition, toxoflavin inhibits growth of *Vermameoba vermiformis*, a host cell of *L. pneumophila*. Our study indicates that *P. alcaliphila* may be important to restrict growth of *L. pneumophila* in water systems through the production of toxoflavin. A sufficiently high concentration is likely not achieved in the bulk water but might have a local inhibitory effect such as in biofilm.

## 1. INTRODUCTION

Legionellosis is a human respiratory disease caused by the bacterium *Legionella* (Cunha et al., 2016). *Legionella pneumophila* cause 90% of the cases; the remaining 10% of cases mostly involve *L. micdadei*, *L. bozemanii*, and *L. Longbeachae* (Cunha et al., 2016). Legionellosis includes Legionnaires’ disease (LD), a systemic infection involving severe pneumonia, and Pontiac fever, a mild, flu-like disease (Cunha et al., 2016). There is a clear upward trend in the prevalence of Legionnaires’ disease worldwide (Cunha et al., 2016). Such an increase in LD cases could be due to densification of urban areas, improvement in diagnostic tests and reporting, a greater number of persons at risk due to aging and/or increase in immunocompromised populations, aging infrastructures, or climate change (Cunha et al., 2016). Importantly, a study of the impact of infectious disease in Europe published in 2018 revealed that LD is the fifth most burdensome disease in people older than 15 years old, after AIDS, tuberculosis, influenza and invasive pneumococcal disease (Cassini et al., 2018).

Not long after the discovery of *Legionella*, it was established that it is transmitted to human by inhalation of aerosols containing *L. pneumophila* that are generated by engineered water systems (EWS) (Meyer, 1983). Several type of EWS can shed *L. pneumophila*, including water distribution systems (showers and faucets), spas, fountains, and cooling towers (Heijnsbergen et al., 2015). Most of the large outbreaks of LD are caused by cooling towers (Garrison et al., 2016). Deficiencies in management and operation of water systems is the main cause of outbreak of LD (Garrison et al., 2016; Mouchtouri et al., 2010).

Cooling towers contain a complex microbial ecosystem constituted by a diverse community of planktonic and biofilm-associated bacteria, protozoa, and algae (Gregorio et al., 2017; Hauer et al., 2016; Llewellyn et al., 2017; Paniagua et al., 2020; Paranjape et al., 2020b, 2020a; Pereira et al., 2017; Pinel et al., 2021; Tsao et al., 2019). The microbial community in cooling towers is mostly found in surface attached and floating biofilms which seeds the bulk water with planktonic microorganisms undefined. The bacterial community identified by different studies varies greatly, but typical biofilm-forming aquatic bacteria were generally identified, such as *Sphingomonadaceae, Caulobacteraceae* and *Hyphomicrobiaceae* (Gregorio et al., 2017; Paniagua et al., 2020; Wang et al., 2013). Local temperature seems to influence the composition of the bacterial and the protozoan community in the attached biofilm (Paniagua et al., 2020). Some protozoa, such as amoeba and ciliates, prey on other microorganisms and *Legionella* has evolved to hijack the endocytic pathway of a variety of these phagocytic protozoans to facilitate their own growth (Boamah et al., 2017; Fields et al., 1984; Paranjape et al., 2020a; Rowbotham, 1980). In fact, most, if not all, of the multiplication of *Legionella spp.* in water system occurs inside these host cells (Boamah et al., 2017). Protozoal hosts also shield *Legionella* against deleterious conditions (Boamah et al., 2017). Therefore, *Legionella*’s presence in engineered water systems is strongly associated with the presence of host cells (Boamah et al., 2017). In multi-species biofilm, *L. pneumophila* has been found associated with amoebas and other phagocytic protozoans and in dense microcolonies (Taylor et al., 2013).

Several bacteria inhibit the growth of L. pneumophila on solid medium, including species of Aeromonas, Bacillus, Flavobacterium, Pseudomonas, Acinetobacter, Kluyvera, Rahnella, Burkholderia, Staphylococcus, Stenotrophomonas or Sphingobacterium (Corre et al., 2021, 2019; Guerrieri et al., 2008; Loiseau et al., 2015; Temmerman et al., 2007; Verdon et al., 2008). The active substances responsible for the inhibition were only identified for a fraction of these species.

*Staphylococcus warneri* produces an antimicrobial peptide named warnericin (Verdon et al., 2008). In the case of *Bacillus*, secreted proteases and surfactin were identified to have an anti-*Legionella* effect (Loiseau et al., 2015). *Pseudomonas fluorescens* produces the volatile compound 1-undecene that inhibits the growth of *L. pneumophila* in a separate but nearby dish, a phenomenon that was called aerial killing (Corre et al., 2021, 2019). These studies indicate that the cooling tower microbiota likely produces a wide range of biomolecules, including proteins, antimicrobial peptides, and biosurfactants, that can have deleterious effects on *L. pneumophila* and its natural hosts in water systems (Berjeaud et al., 2016).

Several studies have shown that the composition of the microbial community in cooling towers is influenced by the disinfection regime (Llewellyn et al., 2017; Paranjape et al., 2020b; Pereira et al., 2017; Tsao et al., 2019). There is also strong inverse relationship between *L. pneumophila* and *Pseudomonas* in cooling towers (Llewellyn et al., 2017; Paranjape et al., 2020b; Tsao et al., 2019). These observations are consistent with our previous study that found that continuous chlorine application reduced microbial diversity, decreasing the abundance of *L. pneumophila* while increasing the abundance of *Pseudomonas* (Paranjape et al., 2020b). *Pseudomonas* is a large genus and several species of *Pseudomonas* inhabit cooling towers (Pereira et al., 2018). Which one correlates negatively with *L. pneumophila* in real cooling towers remains unknown. The goal of the present study was to identify the species of *Pseudomonas* associated with the absence of *L. pneumophila* in cooling towers and identify molecules possibly involved in this competitive relationship.

## 2. MATERIALS AND METHODS

### 2.1 Bacterial strains and cultures

*L. pneumophila* strains Philadelphia 1 (ATCC 33152) and lp120292, which was involved in the LD outbreak in Quebec City in 2012 (Lévesque et al., 2014) and is hereafter referred to as the Quebec strain, were used as test strains. Strains stored at -80°C in 10% glycerol were cultured aerobically at 37°C for 3 days on buffered charcoal yeast extract (BCYE) agar supplemented with 0.25 mg/ml L-cysteine and 0.4 mg/ml ferric pyrophosphate. *Pseudomonas alcaliphila* strain JCM 10630 (CIP 108031T) was acquired from the Centre de Ressources Biologiques de l’Institut Pasteur and grown on nutrient agar at 30 °C. AYE broth (BCYE without agar and charcoal) or Fraquil, an approximate freshwater media (Morel et al., 1975), were used as liquid medium.

### 2.2 *Pseudomonas*-specific 16S amplicon sequencing

The *Pseudomonas*-specific 16S amplicon sequencing strategy previously published by *Pereira et al.* (2018) (Pereira et al., 2018) was used to identify the species of *Pseudomonas* present in 14 cooling towers sampled in a previous study (Paranjape et al., 2020b). A two-step PCR strategy was used to amplify the V3-V4 region of *Pseudomonas* 16S rRNA and add indices using Paq5000 polymerase. The DNA was first amplified with the Pse434F (5’-<u>TCGTCGGCAGCGTCAGATGTGTATAAGAGACAG</u>ACTTTAAGTTGGGAGGAAGGG-3’) and Pse665R (5’-<u>GTCTCGTGGGCTCGGAGATGTGTATAAGAGACAG</u>ACACAGGAAATTCCACCACCC-3’) containing 5’ overhang for Illumina Nextera Indexing kit (underlined). An initial denaturation step of 2 minutes at 95°C was used followed by 30 cycles consisting of 30 s at 95°C, 30 s at 58°C and 30 s at 72°C, and a final elongation step of 7 min at 72°C. The amplicons were purified with AMPure XP beads as per the manufacturer’s instruction. Indexing PCR was then performed with the Nextera XT index kit, according to the manufacturer’s instruction. The amplicons were purified as above and quantified using Quant-iT PicoGreen dsDNA Assay Kit (Invitrogen). The amplicons were sequenced on an Illumina MiSeq using the V2 250 bp paired end reagent kit. The data are available from the Sequence Read Archive under the BioProject accession number PRJNA787128.

The resulting sequences were processed using DADA2 (Callahan et al., 2016) implemented in Qiime 2 version 2018.8 (Bolyen et al., 2019). The sequences were trimmed by 21 nt and truncated to 200 nt. The dataset was rarefied to 50,000 sequences per samples. The taxonomic assignment to the genus level of the resulting amplicon sequence variants (ASV) was assigned using a classifier trained on the SILVA SSU database 132 (Quast et al., 2013), according to Qiime 2 instructions. Species-level taxonomic assignment of the ASVs was performed using BLAST+ against the curated 16S rRNA sequences of *Pseudomonas* as previously described (Pereira et al., 2018). This dataset was then further analyzed using MicrobiomeAnalyst (Chong et al., 2020) to calculate Shannon diversity and perform linear discriminant analysis effect size (LEfSe) (Segata et al., 2011).

### 2.3 *Legionella pneumophila* inhibition assay

*Anti-Legionella* assay was performed using a soft agar overlay technique. Briefly, a suspension of *L. pneumophila* strains, Quebec and Philliladelphia-1, and *P. alcaliphila* were prepared in AYE broth and adjusted to an OD_600nm_ of 0.2. Then 200 µl of *Legionella* suspension was added to 5 ml of autoclaved soft agar (0.5% of agar in ddH2O) and gently poured on the surface of solidified CYE agar plate. The plates were left to solidified in a biological safety cabinet for 15 minutes. Then, a 10 µl drop of *P. alcaliphila* suspension was inoculated in the middle of the plates. Plates were incubated at 25 °C, 30 °C and 37 °C for 3 days and the diameter of the *P. alcaliphila* colony and the zone of inhibition was measured.

### 2.4 Organic extraction of toxoflavin from *P. alcaliphila* supernatant

Chloroform extraction was performed as previously described (Chen et al., 2012) with slight modifications. *P. alcaliphila* was grown on CYE plate for 4 days at 30 °C. The bacterial cells were removed from the surface of the agar using a cell scraper and the agar was cut in smaller pieces using a sterile razorblade. The chopped agar was then mixed with chloroform in 1:1 (w/v) ratio in 50 ml falcon tube for extraction of toxoflavin. The chloroform fraction (∼25 ml) was filtered through Filtropur S 0.2 µm filter (Sarsted) and left to fully evaporated in the fume hood. The extract was then resuspended in 200 µl of methanol. The agar from a sterile CYE plate was processed the same way to serve as a negative control. Extracts were tested with the disc diffusion assay described below.

### 2.5 Disc diffusion assay

The susceptibility of the test strains to commercial toxoflavin (Sigma-Aldrich) and *P. alcaliphila* extracts was assessed by adapting the soft overlay agar technique previously described. Instead of adding the *P. alcaliphila* suspension to the center, a sterile paper disc was placed and 10 ul of extract or a range of commercial toxoflavin concentrations (0 ng/μl, 50 ng/μl, 100 ng/μl, 250 ng/μl and 1000 ng/μl) was added to the disc. Plates were incubated at 30 °C for 3 days and the zone of inhibition was measured.

### 2.6 LC-ESI-MS analysis of toxoflavin

Liquid chromatography/mass spectrometry analysis was performed by using high performance liquid chromatography (HPLC; Waters 2795, Mississauga, ON, Canada) equipped with a 100 × 4.6 mm i.d. Kinetex C8 (Phenomenex) reversed-phase column (particle size 2.6 μm) using a MeCN/H_2_O gradient containing 1% acetic acid at a flow rate of 400 µl/min. The detector was triple quadrupole mass spectrometer (Quattro Premier XE, Waters) equipped with a Z-spray interface using electrospray ionization in positive mode. Analyses were carried out in both MS full-scan and MS/MS multiple reaction monitoring (MRM) scan modes with a mass-to-charge ratio (m/z) window ranging from 130–930. The capillary voltage was set at 3.5 kV and the cone voltage at 30 V. The source temperature was kept at 120 °C and the desolvating gas at 200 °C. Nitrogen was used as the cone and desolvation gas and argon was used as collision gas at collision energies up to 30 eV.

### 2.7 Biofilm and pellicle assay

The biofilm-formation ability of *P. alcaliphila* was investigated by inoculating media with bacterial. Briefly, 1 mL of trypticase soy broth, King’s B, or R2-A broth was added to center four wells of a 24-well plate. Surrounding wells were filled with sterile water to prevent desiccation. Then, 20 µL of *P. alcaliphila* in Fraquil (OD_600nm_ = 0.1) was added to each well and incubated at room temperature with or without shaking at 150 rpm. After a week, images of the wells were taken. When incubated without shaking, plates were shaken at 150 rpm for 1 hour to determine if pellicles could be formed.

### 2.8 *Vermamoeba vermiformis* inhibition assay

The sensitivity of *L. pneumophila* host *Vermamoeba vermiformis* to toxoflavin was determined by monitoring its growth in the presence of toxoflavin. *V. vermiformis* were grown at room temperature in 75 cm^2^ cell culture flasks (Sarstedt) in modified PYNFH medium (ATCC medium 1034) and passaged when confluence was reached. The amoebas were passaged 3 days prior to exposure by adding 5 mL of culture to 20 mL of fresh modified PYNFH. Cell concentration was with a Guava EasyCyte flow cytometer. To prepare samples for flow cytometer, 400 µL of culture was centrifuged at 5000 g for 2 min, the supernatant discarded, and the pellet resuspended in 400 µL of phosphate buffered saline (PBS). The stock culture was diluted to 5 × 10^4^ cells/mL in fresh modified PYNFH and 900 µL was added to the wells of a 24-well plate. Then 100 µL of different toxoflavin solutions were added to wells to give final toxoflavin concentrations of 0, 10, 25, 50 or 100 µg/mL. The plate was incubated at room temperature without shaking. After 2 and 4 days, 400 µL samples were taken from each well to measure cell concentration with flow cytometer. Each condition was performed in triplicate. Results were analyzed using two-way ANOVA, with time and toxoflavin concentration as factors, and Tukey’s test correction for mutltiple comparison was used to access significance between conditions.

## 3. RESULTS

### 3.1 Profiling of *Pseudomonas* species in cooling towers

*Pseudomonas*-specific 16S rRNA amplicon sequencing was performed on triplicate samples from 14 cooling towers. The microbiota of these cooling towers was previosuly studied using 16S rRNA sequencing and 18S rRNA sequencing (Paranjape et al., 2020b, 2020a). For *Pseudomonas*-specific 16S rRNA amplicon sequencing, a total of 4,680,703 reads were obtained and processed with Qiime using the DADA2 pipeline (Bolyen et al., 2019; Callahan et al., 2016). A no template control and DNA extracted from a blank carthridge were also included. As can be seen in Figure 2, these 2 control samples contained very few sequences passing quality control, indicating that the amplicons from the cooling tower samples are not contaminated by spurious sequences. The cooling tower samples contained a minimum of 51089 sequences passing quality control. The data set was rarefied to 50000 sequences per cooling tower. All sequences were assigned to *Pseudomonas*, showing the high specificity of the primers used, as previously reported (Pereira et al., 2018)

**Figure 1:**
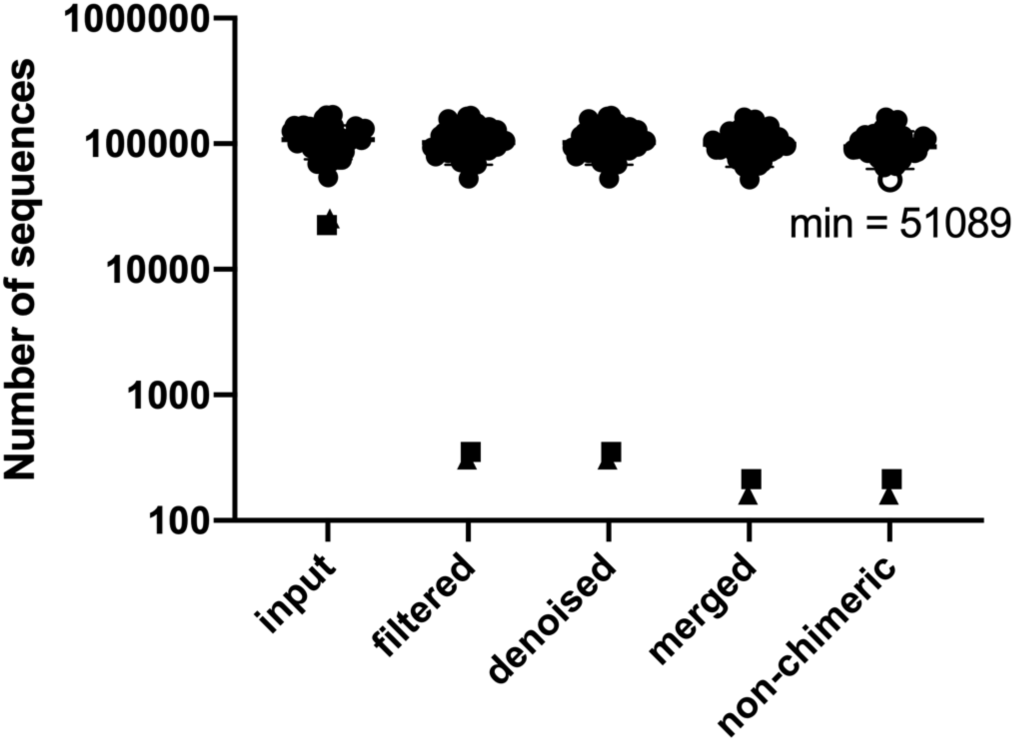
Total number of sequences obtained (input) and left after each one of the processing steps. Square, no template control; triangle, blank carthridge. The sample with the least number of sequences is depicted by an open circle.

**Figure 2.**
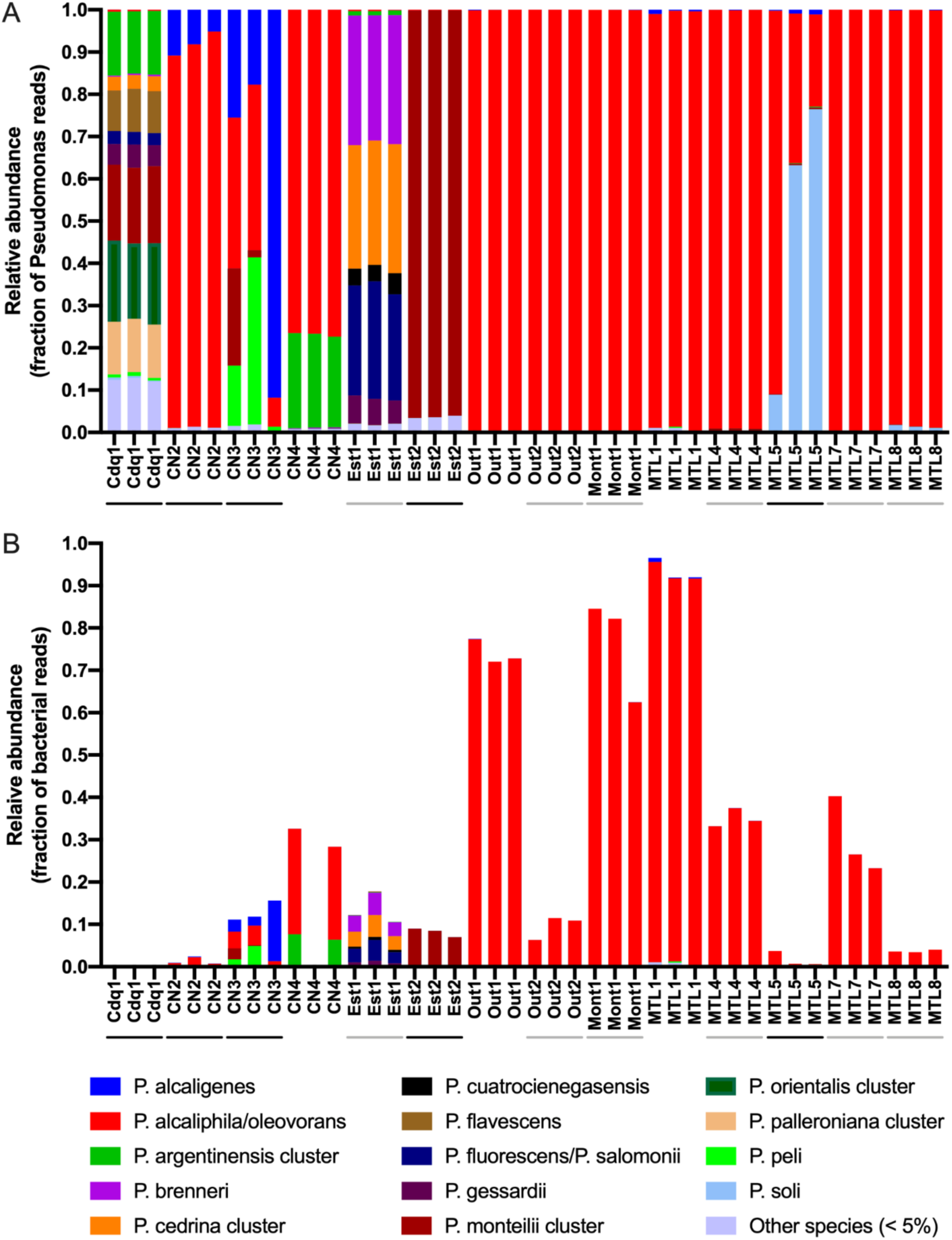
Relative abundance of *Pseudomonas* species as a fraction of *Pseudomonas* reads (A) and as a fraction of bacterial reads (B). Species with a maximum relative abundance of less than 5% in any towers were grouped together in the category “other species”. The presence of *Legionella* in each tower is depicted by a line under the cooling tower’s name; a grey line indicate the presence of *Legionella* species other than *pneumophila*, a black line indicate the presence of *L. pneumophila* (Paranjape et al., 2020b).

Considering all the cooling towers, 34 *Pseudomonas* species were found among which 14 can be described as major or abundant species and the other 20 as minor species, collectively representing less than 5% of the population. Of note, this method is unable to differentiate closely related species, such as *P. alcaliphila* and *P. oleovorans*. Such species are therefore grouped in clusters. The highest diversity of *Pseudomonas* species was observed in cooling towers Cdq1 and Est1 containing 26 and 13 different species, respectively, followed by CT CN3, CN4 and MTL5 (Figure 2A). Gobally, the top three most abundant *Pseudomonas* species in the studied cooling towers were *P. alcaliphila/oleovorans*, *P. monteilii* and *P. alcaligenes. P. alcaliphila/oleovorans* was observed in nearly 100% of cooling towers in various proportions, but was the largely dominant *Pseudomonas* species in several cooling towers including CN2, Out1, Out2, Mont1, MTL1, MTL4, MTL7 and MTL8 (Figure 2A). The human pathogen *P. aeruginosa* was detected only in towers CdQ1 and Est2 at a low abundance of 0.01. Next, we calculated the abundance of each *Pseudomonas* species as a fraction of relative bacterial abundance (Paranjape et al., 2020b). As can be seen in Figure 2B, *P. alcaliphila/oleovorans* is the most abundant *Pseudomonas* species in cooling tower microbiomes dominated by *Pseudomonas*, including cooling towers Out1, Mont1, and MTL1.

Next, we sought to identify factors influencing the diversity of *Pseudomonas* within the cooling towers. Shannon diversity was calculated for each tower (Figure 3A). As expected, cooling towers CdQ1 and Est1 have the highest shannon diversity, while the cooling towers dominated by *P. alcaliphila* had the lowest diversity. In our previous study, we determined that chlorine application schedule had a greater impact than chlorine concentration in shaping the bacterial communities (Paranjape et al., 2020b). Similarly, treatment schedule seems to affect *Pseudomonas* diversity as tower treated by periodic applications showed significantly less diversity (*P* = 0.03) than cooling towers treated continuously (Figure 3B). Concentration of total chlorine and residual chlorine did not seem to affect *Pseudomonas* diversity. The slopes of the Shanon Diveristy against the concentration of total chlorine (−0.599 ± 0.502) and against the concentration of residual chlorine (−0.107 ± 0.073) were not significantly different than zero (Figure 3C and 3D).

**Figure 3:**
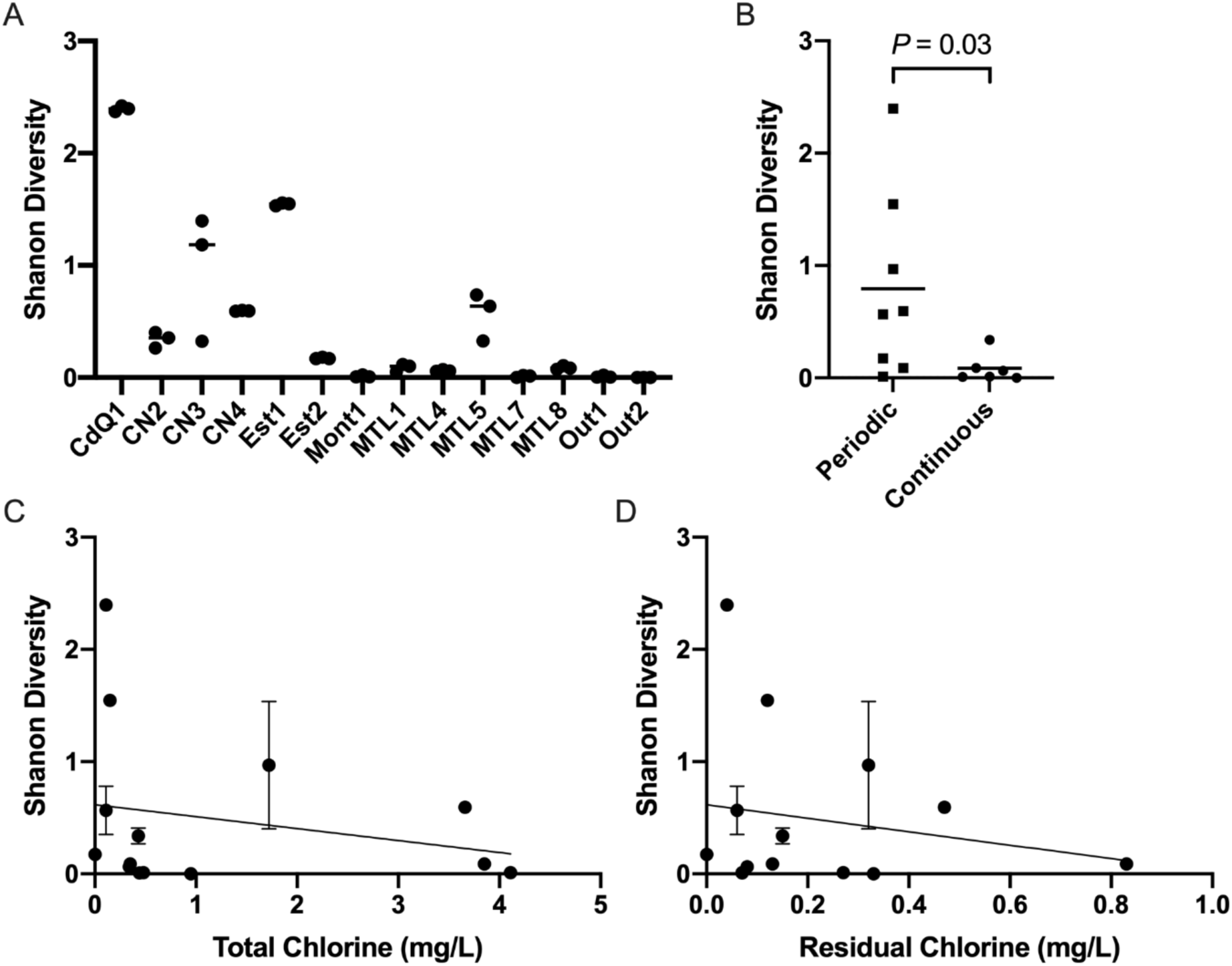
Treatment application schedule but not chlorine concentration affect *Pseudomonas* diversity. Shanon diversity index was calculated for each cooling towers (A) to evaluate alpha diversity. Individual replicate values are shown. Average shanon diversity index was calculated and each tower was classified according to the treatement schedule (B). A one-tail t-test was used to access statistical significance between the two groups. The average Shanon diversity index ± standard deviation was plotted against total chlorine (C) and redidual chlorine concentration (D). A simple linear regression model was used to determine if the slope was significantly different than zero.

We next used linear discriminant analysis effect size (LEfSe) implemented in MicrobiomeAnalyst (Chong et al., 2020) to examine the differences in the abundance of *Pseudomonas* species relative to the bacterial reads in cooling towers. LEfSe is an algorithm that uses a mix of statistical testing, linear discriminant analysis (LDA), and effect size to identify taxa that are predictive of a particular condition (Segata et al., 2011). The treatment schedule and the presence or absence of *Legionella spp.* and *Legionella pneumophila* were considered as comparison factors (Paranjape et al., 2020b). *P. alcaliphila/oleovorans* was the only species enriched in cooling towers free of *L. pneumophila* and free of *Legionella spp.* (Figure 4A and B). In contrast, *P. montelli* cluster was enriched in towers containing *Legionella* or *L. pneumophila*, *P. alcaligenes* was enriched in cooling towers containing *L. pneumophila,* and *P. soli* was enriched in towers containing *Legionella spp.* (Figure 4A and B). By considering the treatment schedule, *P. alcaliphila/oleovorans* and *P. anguiliseptica* were associated with continuous treatment whereas *P. monteilii* and *P. soli* were associated with periodic treatment (Figure 4C).

**Figure 4.**
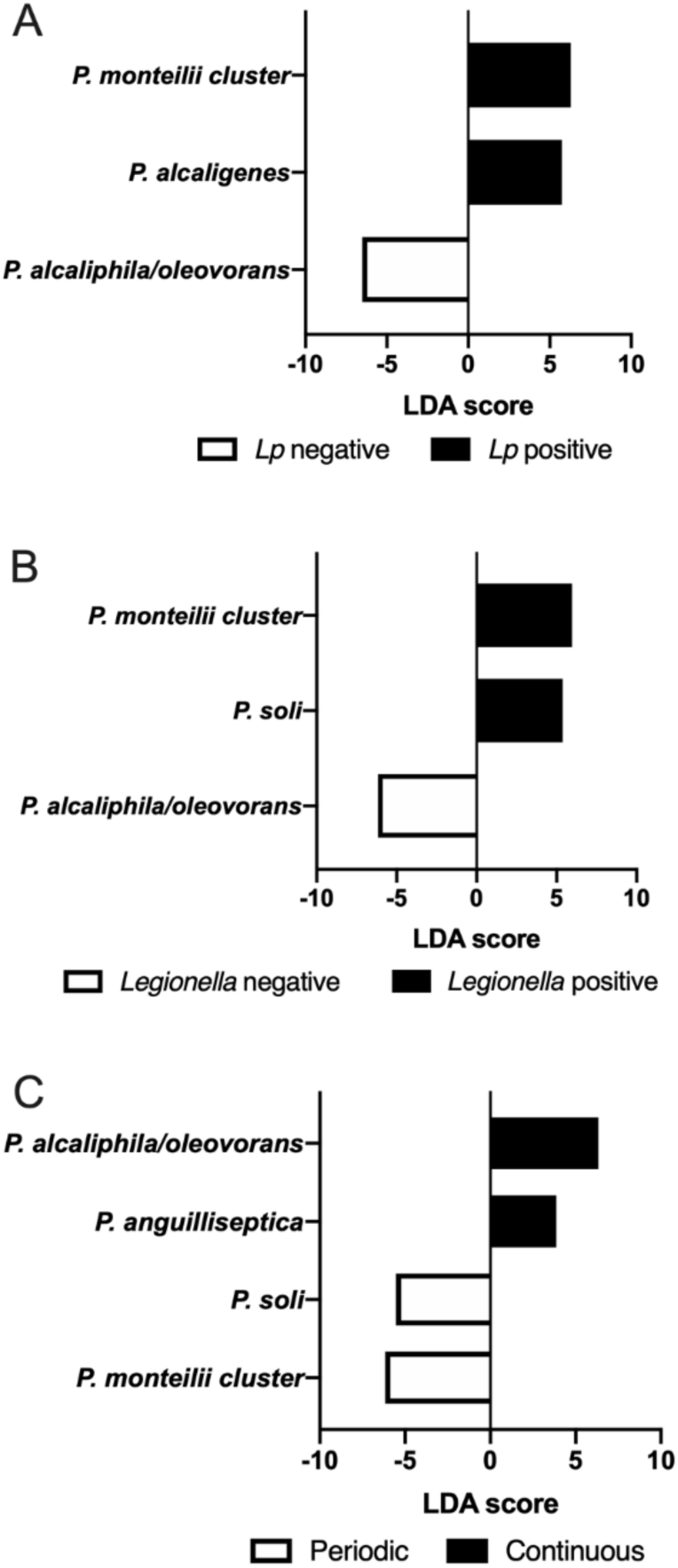
*P. alcaliphila/oleovorans* is enriched in towers free of *Legionella* and associated with continuous treatement. LDA scores of *Pseudomonas* species associated with the presence of *L. pneumophila* (A), the presence of *Legionella spp.* (B) and treatment schedule (C). Only species with *P* < 0.05 are shown.

### 3.2 Anti-*L. pneumophila* activity of *P. alcaliphila*

Our results reavealed that a member of the *P. alcaliphila/oleovorans* cluster seems to be the main inhibitor of *L. pneumophila* colonisation in the cooling towers we studied. Therefore, we investigated if an isolate of that cluster, *P. alcaliphila* strain JCM 10630, can inhibit *L. pneumophila* growth *in vitro* (Yumoto et al., 2001).

We carried out *L. pneumophila* inihibition assay at three different temperatures: 25 °C, 30 °C and 37 °C. After three days of incubation, *P. alcaliphila* inhibited the growth of *L. pneumophila* at 25 °C and 30 °C but not at 37 °C (Figure 5). A one-way ANOVA with a Tukey correction for mutltiple comparison was used to access significance between conditions. The diameter of inhibition was significantly larger at 25 °C than at 30 °C for both strains (*P* < 0.001). The two strain tested showed similar inhibition zones at each temperature tested (*P >* 0.6). The size of the colony of *P. alcaliphila* was significantly larger at 25 °C than at 30 °C (*P* < 0.001). The strain of *L. pneumophila* seems to influence slighlty the growth of *P. alcaliphila* at 25 °C as the colony was slightly larger (14.3 mm) when grown with the Quebce strain than with the Philadelphia-1 strain (13 mm, *P* = 0.02). There were no difference in colony size at 30 °C.

**Figure 5.**
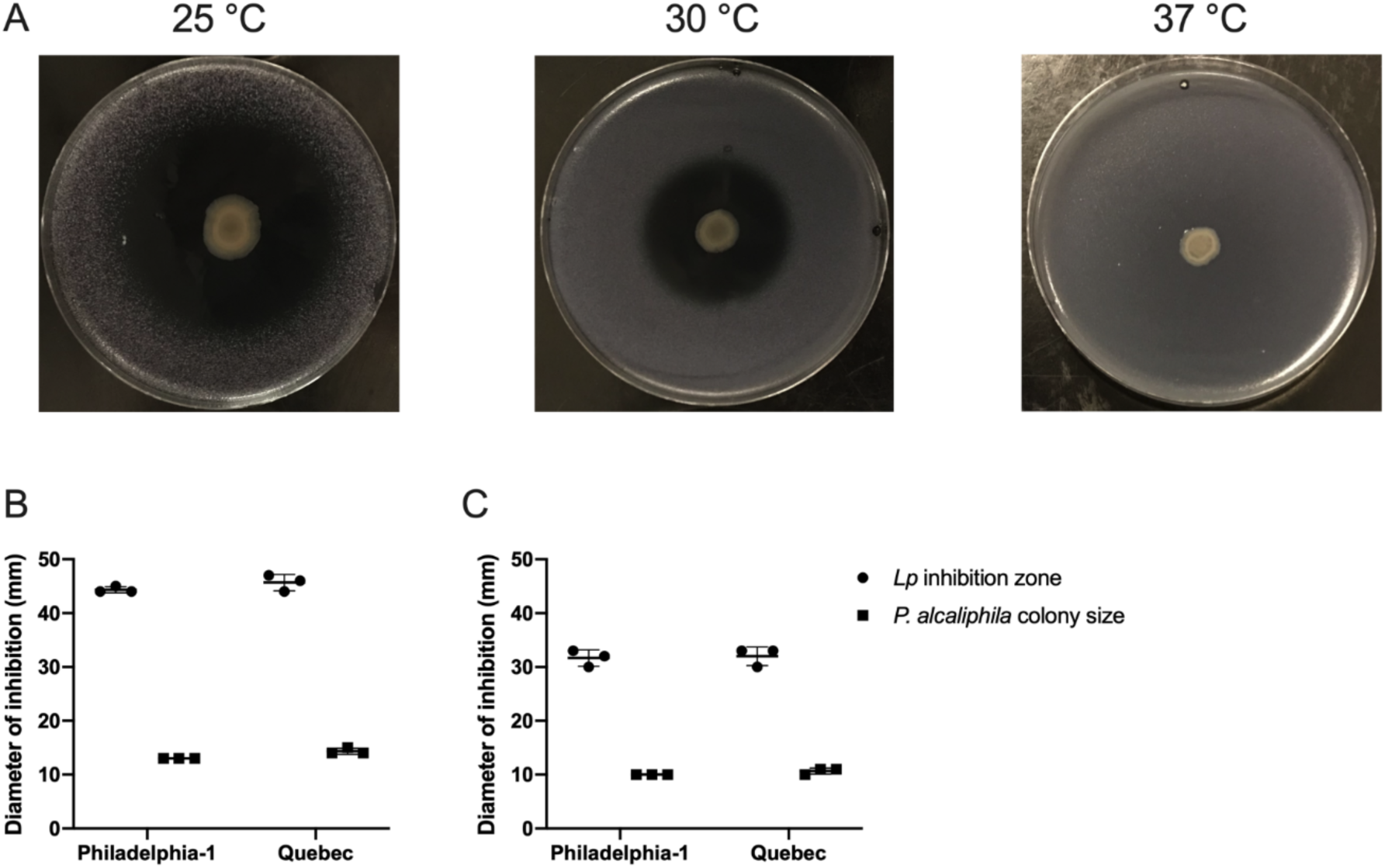
*P. alcaliphila* inhibits the growth of *L. pneumophila* at 25 °C and 30 °C. *L. pneumophila* starin Philadelphia-1 and the Quebec strain were inoculated on CYE plates in a thin layer of soft agar. Once solidified, *P. alcaliphila* was inoculated in the center of the plate. A) Representative image of plates incubated at 25 °C (left panel), 30 °C (center panel) and 37 °C (right panel). The diameter of inhibition (circle) and the diameter of the *P. alcaliphila* colony (square) for three replicates was recorded for plates incubate at 25 °C (B) and at 30 °C (C).

### 3.3 In silico analysis of P. alcaliphila genome

The *P. alcaliphila* strain JCM 10630 genome was retrieved from RefSeq (GCF_900101755.1) and was analysed to identify clues as to the cause of the inhibition of *L. pneumophila* growth. We first used antiSMASH (Blin et al., 2019) to identify putative biosynthetic gene clusters (BCGs). Five clusters were found but none showed similarity higher than 50% with known clusters (Table 1). Next, we used the Blast KOALA function of the Kyoto Encyclopedia of Genes and Genomes to assign Kegg orthology annotation to the genes and predict metabolic pathways present in this genome (Kanehisa et al., 2016). A cluster of genes homologous to toxoflavin synthesis cluster was detected. Toxoflavin is an improtant virulence factor of the plant pathogen *B. glumae* (Suzuki et al., 2004). Toxoflavin is also produced by *Pseudomonas protegens* Pf-5 (Philmus et al., 2015). The *P. alcaliphila* toxoflavin cluster is most homologous to *B. glumae* cluster and organized in a similar manner (Chen et al., 2012; Philmus et al., 2015; Suzuki et al., 2004). The homology of *P. alcaliphila* genes compared to *B. glumae* varries between 69% identity to 36% identity (Table 2). Our *in silico* analysis suggests that the inhibition of *L. pneumophila* growth by *P. alcaliphila* could be due to the production of toxoflavin, another compound, or a mixture of several molecules.

**Table 1.**
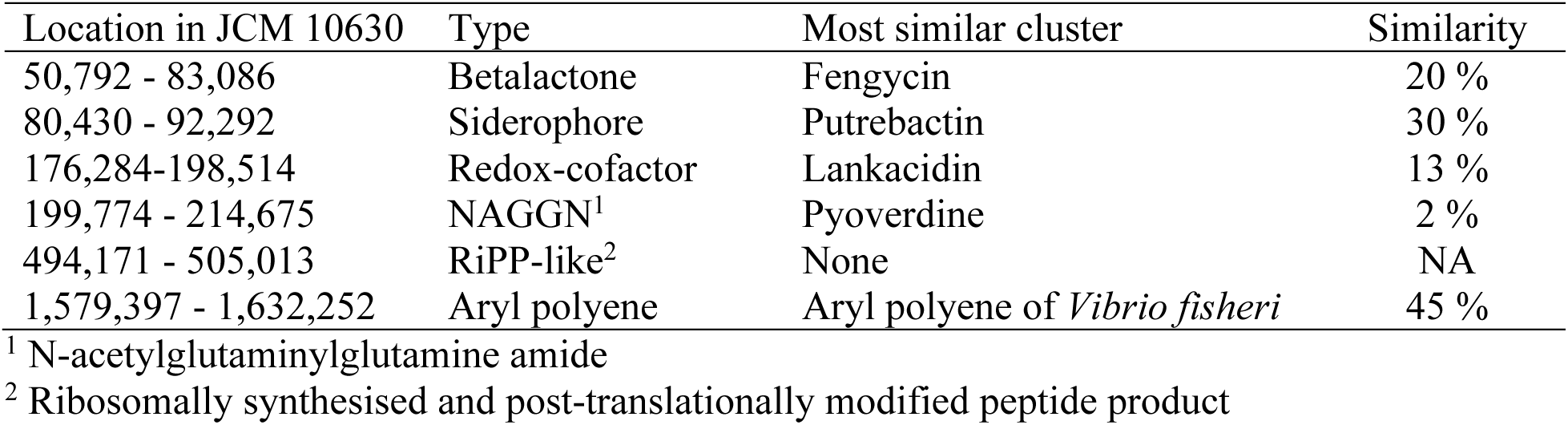
Biosynthetic gene clusters identified in the genome of *P. alcaliphila* JCM 10630 with antiSMASH.

**Table 2.** Comparison of the genes in *P. alcaliphila* toxoflavin cluster with *B. glumae* homologues.

homologues
| Locus tag | Protein product | Length | Annotated function | Name | *B. glumae* homologue (% identity) |
| --- | --- | --- | --- | --- | --- |
| PAL02S_RS12525 | WP_074675559.1 | 388 | bifunctional diaminohydroxyphosphoribosylaminopyrimidine deaminase/5-amino-6-(5-phosphoribosylamino)uracil reductase | ToxE | bglu_2g06440 (42%) |
| PAL02S_RS12530 | WP_074675558.1 | 562 | WD40 repeat domain-containing protein | ToxC | bglu_2g06420 (47%) |
| PAL02S_RS12535 | WP_074675557.1 | 201 | GTP cyclohydrolase II | ToxB | bglu_2g06410 (44%) |
| PAL02S_RS12540 | WP_139202874.1 | 340 | SUMF1/EgtB/PvdO family nonheme iron enzyme | ToxD | bglu_2g06430 (54%) |
| PAL02S_RS12545 | WP_074675555.1 | 245 | class I SAM-dependent methyltransferase | ToxA | bglu_2g06400 (68%) |
| PAL02S_RS12550 | WP_074675554.1 | 302 | LysR family transcriptional regulator | ToxR | bglu_2g06390 (69%) |
| PAL02S_RS12555 | WP_074675553.1 | 157 | DMT family transporter | ToxF | bglu_2g06380 (64%) |
| PAL02S_RS12560 | WP_074675552.1 | 364 | efflux RND transporter periplasmic adaptor subunit | ToxG | bglu_2g06370 (36%) |
| PAL02S_RS12565 | WP_074675551.1 | 1023 | MexW/MexI family multidrug efflux RND transporter permease subunit | ToxH | bglu_2g06360 (55%) |

### 3.4 Toxoflavin inhibits growth of *L. pneumophila* on CYE plate

The susceptibility of the two strains of *L. pneumophila* to toxoflavin was tested using a dilution series of commercial toxoflavin (Sigma-Aldrich). The results showed that the size of the inhibition zone proportionally increases with concentration of toxoflavin for both strains (Figure 6). The diameters of the inhibition zone were 10 mm, 12 mm, 20 mm and 30 mm for concentrations of 0.5, 1, 2.5 and 10 µg, respectively.

**Figure 6.**
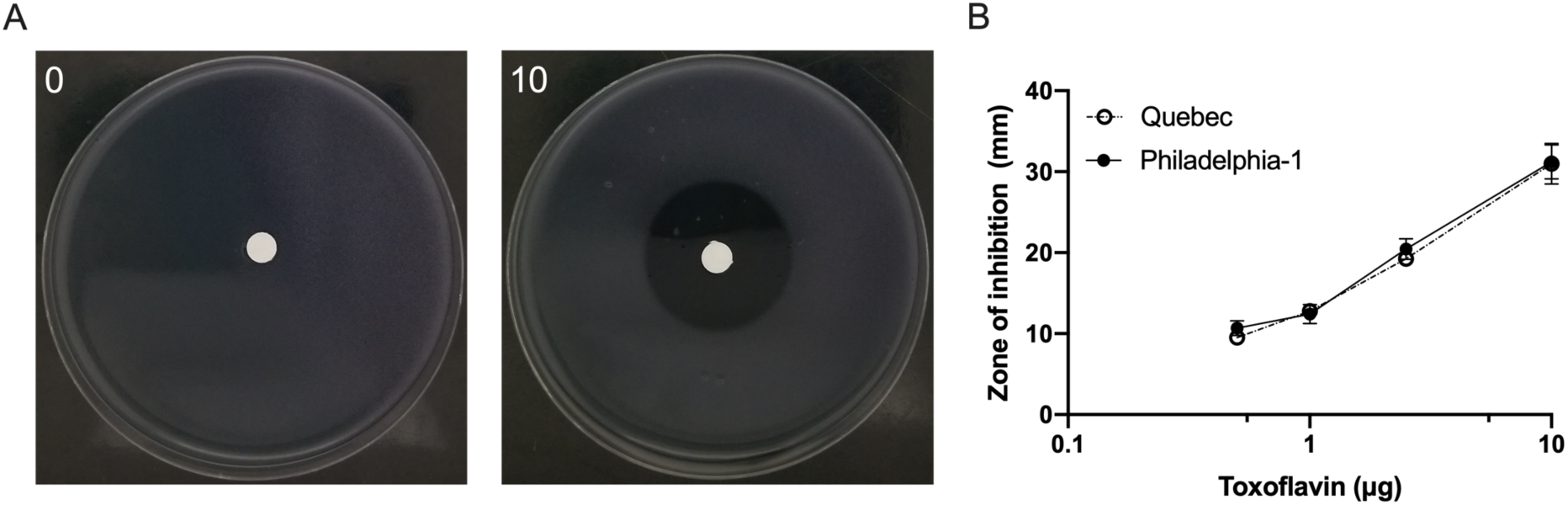
Toxoflavin inhibits the growth of *L. pneumophila* on CYE plates. A disc diffusion assay was used to test the susceptibility of two *L. pneumophila* strains to 0.5, 1, 2.5 and 10 µg of toxoflavin at 30°C. Representative image for *L. pneumophila* strain Philadelphia-1 at 0 and 10 µg is shown in A. The average and standard deviation of the size of the zone of inhibition for each concentration tested in triplicate for *L. pneumophila* strain Quebec and Philadelphia-1 is shown in B.

### 3.5 Toxoflavin is secreted by *P. alcaliphila* on CYE agar plate

In order to confirm that *P. alcaliphila* produces toxoflavin, we performed chloroform extraction from CYE plate inoculated with a pure culture of *P. alcaliphila.* Controls included an extract from a sterile CYE plate and the methanol carrier alone. *L. pneumophila* growth was inhibited by the extract from plates inoculated with *P. alcaliphila* (Figure 7C) slightly more than sterile CYE and methanol (Figure 7A and B), with zone of inhibitions of 12, 10 and 10 mm, respectively. In order to confirm that toxoflavin was present, the extracts were then subjected to LC-ESI/MS. Pure toxoflavin solution produced a strong peak at m/z= 194.0 (Figure 7D). The same strong peak appeared in extract from *P. alcaliphila* plate extracts (Figure 7F) while being absent in the control (Figure 7E). These results were also confirmed with LC-ESI-MS/MS in MRM mode (Supplementary Figure S1).

**Figure 7.**
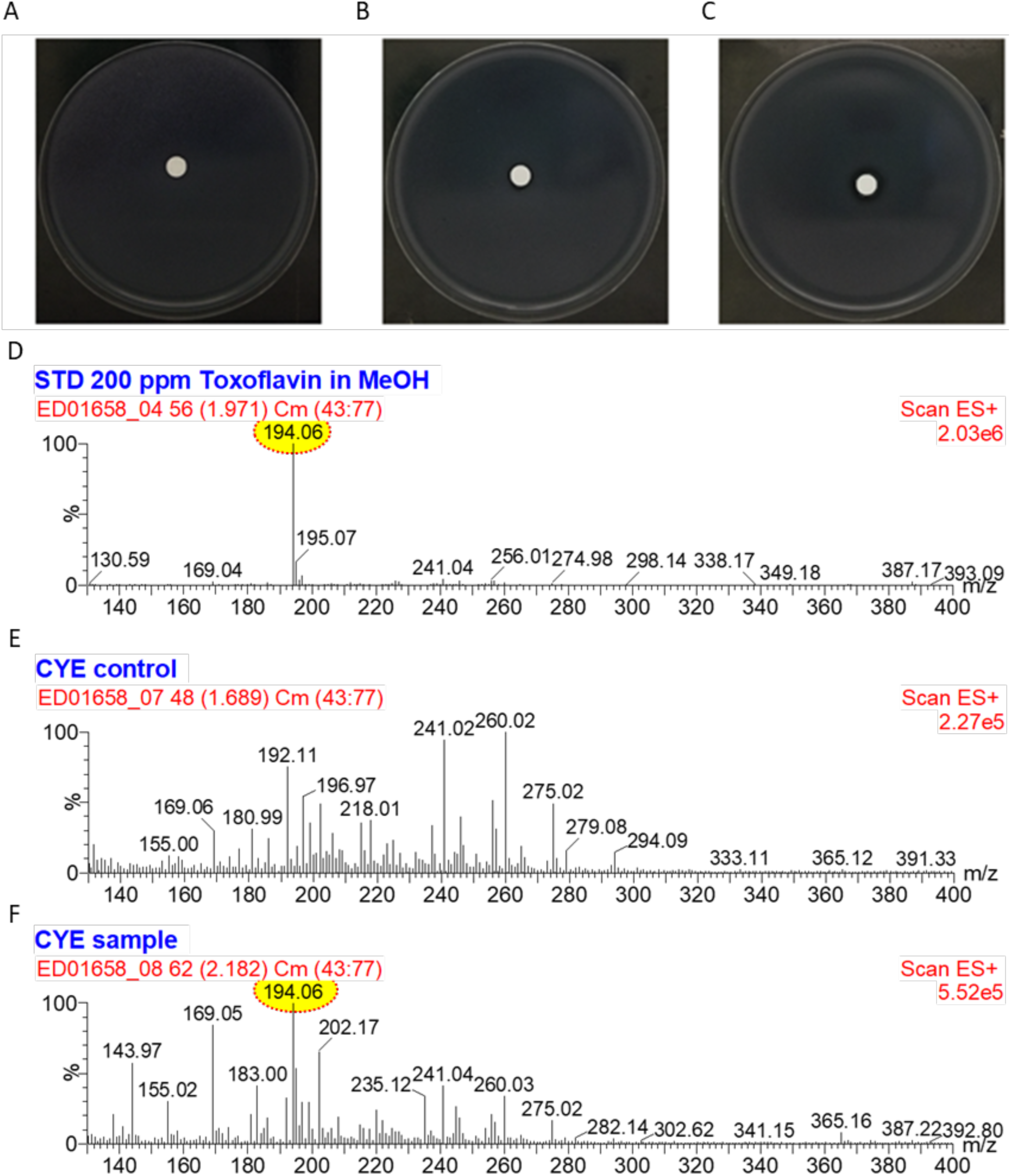
Organic extract of *P. alcaliphila* culture inhibits *L. pneumophila* growth and contains toxoflavin. Disc diffusion assay was used to test the growth inhibition activity of methanol (A), organic extracts from sterile CYE plate (B), and CYE plate covered by a lawn of *P. alcaliphila* (C). The production of toxoflavin (m/z = 194.0) by *P. alcaliphila* was confirmed with LC-ESI-MS by analysing the mass spectrum of pure toxoflavin (D) and comparing it to those of extract from sterile CYE plate (E) and extract from CYE plate covered by *P. alcaliphila* (F).

### 3.6 *P. alcaliphila* produces floating biofilm mat

Many *Pseudomonas* species can produce attached biofilms or floating biofilm mats (Mann and Wozniak, 2012). Therefore, we investigated the ability of *P. alcaliphila* JCM 10630 to form these structures. First, we tested the production of attached biofilm in R2A, King’s B, and in trypticase soy broth at room temperature under shaking. After one week, no attached biofilm was seen, however a filamentous floating mass of cells could be seen in both R2A and trypticase soy broth (Figure 8A). We then tested the production of a pellicle by incubating *P. alcaliphila* in the same three media but without shaking. No pellicle was formed in any of these media (Figure 8B). Nevertheless, we could see a mat at the bottom of the well in all case. Shaking for 1 h dislodge the mat produced in trypticase soy broth and R2A, resulting in a floating mat similar to what was seen after incubation with shaking (Figure 8A). We therefore concluded that *P. alcaliphila* may produces floating biofilm mats in cooling towers.

**Figure 8.**
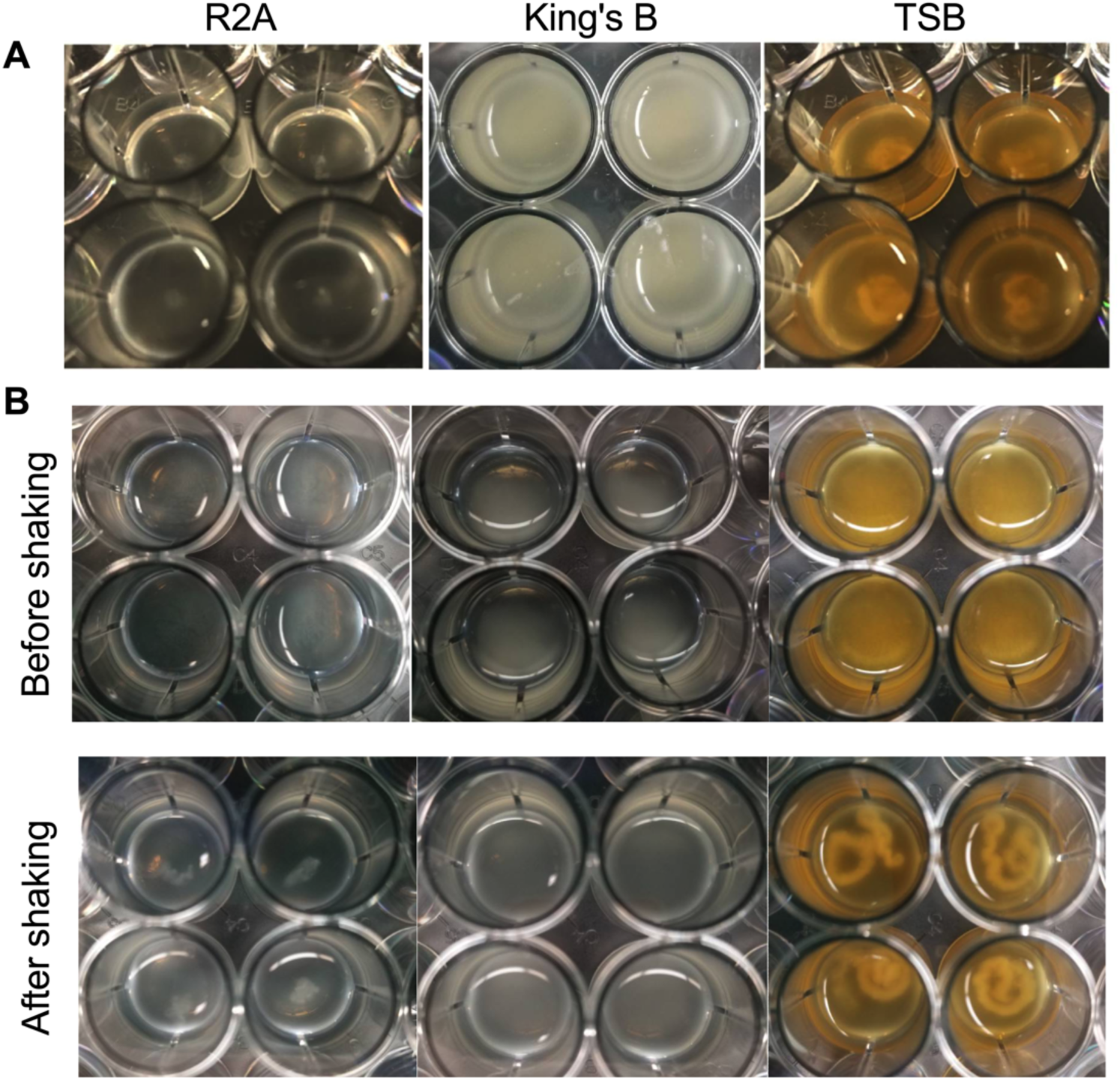
*P. alcaliphila* produced floating biofilm mat. A) R2A, King’s B and trypticase soy broth were inoculated with *P. alcaliphila* and incubated at room temperature with shaking (150 rpm). B) R2A, King’s B and trypticase soy broth was inoculated with *P. alcaliphila* and incubated at room temperature without shaking. After incubation, plates were shaken (150 rpm) for 1h. Representative images of 4 wells of a 24-well plates are shown.

### 3.7 Toxoflavin is toxic for *Vermamoeba vermiformis*

Since *P. alcaliphila* was also negatively correlated with the presence of host cells in cooling towers (Paranjape et al., 2020a), we next hypothesize that toxoflavin might be toxic for amoebas typically found in water systems. Therefore, we monitored the growth of *V. vermiformis* when exposed to toxoflavin (Figure 9). Within four days, cells unexposed to toxoflavin grew by 7.6-fold. In contrast, cells exposed to 10 µg/ml and 25 µg/ml, grew much less, by a factor of 3.7 and 3.3 respectively. Cells exposed to higher concentration show a sharp decrease in cell number at day 2. By day 4 the cells have grown back to the number of cells present at the start of the experiment, suggesting that toxoflavin is metabolized by *V. vermiformis* over time.

**Figure 9:**
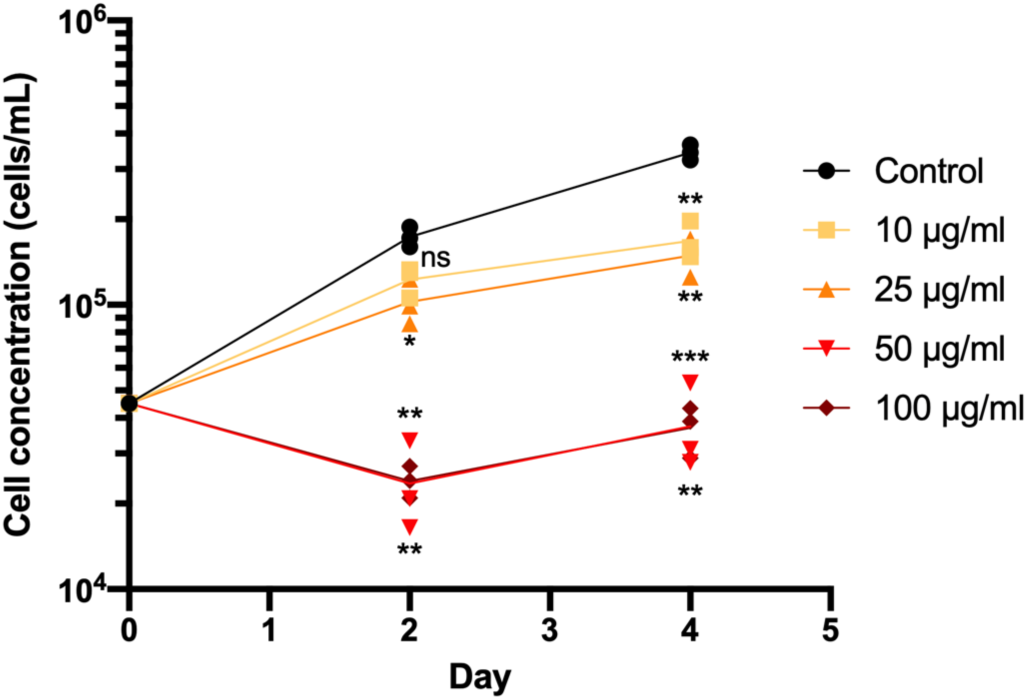
Toxoflavin inhibits axenic growth of *V. vermiformis*. *V. vermiformis* was cultured in PYFNH to a concentration of 50,000 cells/mL and exposed to 10 µg/ml, 25 µg/ml, 50 µg/ml and 100 µg/ml of toxoflavin. Cultures without toxoflavin served as a control. Cell concentration was determined using flow cytometry at day 0, 2 and 4. A two-way ANOVA with a Tukey correction for mutltiple comparison was used to access significance of each test conditions compared to the control (ns, non-significant, * *P* < 0.05, ** *P* < 0.01, *** *P* < 0.001)

## 4. DISCUSSION

Several previous studies have shown an inverse relationship between the presence of *L. pneumophila* and *Pseudomonas* in cooling towers (Llewellyn et al., 2017; Paranjape et al., 2020b). By using *Pseudomonas*-specific 16S rRNA amplicon sequencing approach (Pereira et al., 2018), we identified that members of *P. alcaliphila/oleovoran* cluster are the main species associated with the exclusion of *L. pneumophila* from cooling towers. The dominance of *P. alcaliphila/oleovorans* is surprising as a previous study of a single cooling towers located in Braunschweig, Germany, found a high diversity of species even when *P.alcaliphila/oleovorans* was a member of the core community (Pereira et al., 2018). Differences in source water or cooling tower management could explain the dominance of *P. alcaliphila/oleovorans* in our cooling tower samples compared to those in Germany. The diversity and abundance of *Pseudomonas* may also be affected by seasons, but this could not be investigated with our current data set. Importantly, the presence of *P. alcaliphila/oleovorans* was strongly associated with continuous disinfectant application. This strategy presumably creates conditions favorable for this group or specific members of this group.

Our analysis also revealed that some species of *Pseudomonas* may be beneficial for *L. pneumophila*. LEfSe analysis suggest that members of the *P. monteilii* cluster and *P. alcaligenes* are positively associated with *L. pneumophila* (Figure 4A). For example, it has been reported that in an environment lacking critical nutrients for its growth, *L. pneumophila* can form microcolonies around certain aquatic bacteria including *Flavobacterium breve* (Wadowsky and Yee, 1983) and *Pseudomonas alcaligenes* (Çotuk et al., 2005).

Unfortunately, the method used is not able to differentiate between members of the *P. alcaliphila/oleovorans* cluster as the region targeted is identical. In addition to *P. alcaliphila* and *P. oleovorans*, this cluster also contain *P. chengduensis* (Pereira et al., 2018). These three species are associated with various water environment (Peix et al., 2018; Tao et al., 2014). It is possible that the cooling towers studied here contain a diversity of species belonging to this cluster. The water of the cooling towers included in this study was typically between 20-25 °C and pH 8 (Paranjape et al., 2020b). This falls within the conditions that *P. alcaliphila* JCM 10630 can thrive in, having been isolated from sea water and shown to be alkali-tolerant and psychrophilic, growing best at temperature between 4 and 30 °C (Yumoto et al., 2001).

In this study, we found that *P. alcaliphila* was able to inhibit *L. pneumophila* growth was, at least in part, through toxoflavin production. We found that 0.5 µg of toxoflavin directly inhibit *L. pneumophila* growth on plates, and that a concentration of 25 µg/mL inhibits the growth of *L. pneumophila* host *V. vermiformis.* Genomic analysis revealed that *P. alcaliphila* also contains a homologue of the toxoflavin biosynthetic cluster and the presence of toxoflavin was confirmed in corresponding organic extracts. Inhibition appeared to be temperature-dependent since *P. alcaliphila* inhibited *L. pneumohila* growth at 30°C but not 37 °C. Multiple mechanisms could explain this phenomenon. It could be that the rate of growth of *L. pneumophila* is far greater than *P. alcaliphila* at 37°C, and so outpaced the accumulation of toxoflavin. The other possibility is that temperature affects toxoflavin production. To our knowledge, regulation of toxoflavin production by temperature has not been reported in *Pseudomonas spp*. However, *Pseudomonas* is recognized for having complex quorum sensing systems, which have been studied extensively in *P. aeruginosa* (Chadha et al., 2021). Therefore, the regulation of toxoflavin production by quorum sensing could explain this result. The toxoflavin biosynthetic gene cluster has been thoroughly studied in detail in *B. glumae* and in *Pseudomonas protegens* (Chen et al., 2012). In *B. glumae*, toxoflavin is regulated by quorum sensing system involving TofI, encoding the *N*-octanoyl homoserine lactone synthase, and the associate receptor TofR (Chen et al., 2012; Kim et al., 2004). In turn, TofR induced expression of ToxJ, which induced expression of ToxR, the main regulator of the toxoflavin biosynthesis and transporter loci (Chen et al., 2012; Kim et al., 2004). The *P. alcaliphila* JCM 10630 genome contains a homologue of tofR (40% identity) but no homologue of TofI could be identified. Whether or not toxoflavin production is regulated by quorum sensing system in *P. alcaliphila* will require additional experiments.

It is extremely unlikely that *P. alcaliphila* can produce enough toxoflavin for it to accumulate in cooling towers and reach inhibitory concentrations in bulk water. However, local concentrations in biofilms or floating mats could potentially reach inhibitory or lethal concentrations for *L. pneumophila* and/or *V. vermiformis* since *Pseudomonas* are known to form a diversity of biofilm structure (Koza et al., 2020). Indeed, we showed that *P. alcaliphila* cells can aggregate in pure culture to form floating mats, but not an attached biofilm. This does not eliminate the possibility that *P. alcaliphila* can colonize surfaces in water system or multispecies biofilm. We also cannot rule out that other compounds contributed to the inhibition of *L. pneumophila* by *P. alcaliphila*. Of note, RiPP-like type biosynthetic gene clusters includes bacteriocin and other antimicrobial peptide-derived molecules (Arnison et al., 2012). Similarly, it is possible that the member of the *P. alacaliphila/oleovorans* cluster found in cooling towers is different from the isolate used in this study, and so other or additional BCG and corresponding molecules could be acting as *L. pneumophila* inhibitors.

## 5. CONCLUSION

*Pseudonomas* specific 16S amplicon sequencing increased the species-level resolution of our data which allowed us to narrow down candidate species responsible for the exclusion of *L. pneumophila* in cooling towers. The absence of *L. pneumophila* was most strongly associated with the presence of *P. alcaliphila/oleovorans*. Inspection of the genome of a member of *P. alcaliphila/oleovorans* revealed a biosynthetic gene clusters (BCGs) homologous to toxoflavin synthesis cluster of *B. glumae*. Using LC-MS/MS, toxoflavin was found in extracts from *P. alcaliphila* agar. Toxoflavin is known to be toxic to many microorganisms, we confirmed that this is also the case for both *L. pneumophila* and its host *V. vermiformis* using commercial toxoflavin. To our knowledge, this is the first report of a *P. alcaliphila* strain that produces toxoflavin and which inhibits the growth of *L. pneumophila*. In water systems, this inhibitory effect would be strongest near or within densely populated biofilms or floating mats that *P. alcaliphila* is capable of forming in different media. Finally, toxoflavin was only one of many candidate molecules, as other biosynthetic gene clusters were identified in *P. alcaliphila*. Multiple molecules could be contributing to inhibitory effects of *Pseudomonas* in cooling towers and will need to be investigated.

## ACKNOWLEDGEMENT

We are indebted to Rui P. A. Pereira for sharing the *Pseudomonas* phylogeny database and providing guidance on performing *Pseudomonas*-specific 16S amplicon sequencing. We are thankful to Jesse Shapiro and his team for assistance regarding analysis of the amplicon sequencing data. This work was supported by a FRQNT Team grant (2016-PR-188813) and a Natural Science and Engineering Research Council of Canada Discovery Grant (RGPIN/04499-2018) to SPF. The graphical abstract was created with BioRender.com.

